# Position-dependent effects of *SCN2A* premature stop codons on neuronal excitability and behavior

**DOI:** 10.1101/2025.10.30.685422

**Authors:** Ahmad Al Saneh, Maria Fernanda Hermosillo Arrieta, Lionel Gissot, Madison O’Bryan, Rui Li, Gordon Buchanan, Christopher A. Ahern, Aislinn J. Williams

## Abstract

*SCN2A* encodes the voltage-gated sodium channel Na_V_1.2, a key determinant of spike initiation and propagation in glutamatergic neurons. Premature termination codons are often assumed to produce uniform haploinsufficiency via nonsense-mediated decay, yet whether distinct *SCN2A* premature stop codons yield equivalent molecular, cellular, and behavioral outcomes remains unknown.

We generated two mouse lines carrying patient mutations—*Scn2a^Y84X/+^* (p.Tyr84UAA; early coding sequence) and *Scn2a^R1627X/+^*(p.Arg1627UGA; terminal coding exon)—on a C57BL/6J background. Allele-specific expression was quantified by targeted next-generation sequencing of whole-brain reverse transcribed cDNA. Na_V_1.2 protein was measured in half-brain lysates by automated western blot and *ex vivo* whole-cell recordings were obtained from layer 5b pyramidal-tract neurons in medial prefrontal cortex. A panel of behavioral assays assessed locomotion/exploration, motor learning, anxiety-like behavior, sociability, sensorimotor gating, and seizure susceptibility.

Allele-specific RNA handling diverged by position: mRNA carrying Y84X engaged partial nonsense-mediated decay, whereas R1627X transcripts were at allelic balance. Despite this difference in RNA fate, Na_V_1.2 protein was comparably reduced in both lines. Electrophysiologically, both premature termination codon mutations slowed the action-potential upstroke, with a larger decrement in *Scn2a^Y84X/+^*than in *Scn2a^R1627X/+^*. Spike threshold was depolarized only in *Scn2a^Y84X/+^*, whereas *Scn2a^R1627X/+^* remained similar to wild type. Frequency–current relations showed reduced firing at near-rheobase inputs in both mutants, with responses approaching wild type at stronger currents. Behaviorally, locomotion, sociability, and sensorimotor gating were preserved. Both lines exhibited increased grooming—consistent with restrictive, repetitive behavior; *Scn2a^Y84X/+^*alone showed greater exploration in the elevated-risk context and a male-predominant deficit in rotarod learning. In maximal electroshock testing, mortality was lower in both lines without differences in seizure threshold or severity.

Our results show that distinct *SCN2A* premature termination codons are not equivalent to one another, nor to a uniform haploinsufficient state. An early, nonsense-mediated decay-competent premature stop codon (Y84X) and a terminal-exon one (R1627X) produce partially overlapping yet allele-specific effects on neuronal excitability and behavior. These findings establish premature termination codon position as a determinant of phenotype, supporting allele-tailored mechanistic studies and therapeutic strategies.

## Introduction

Voltage-gated sodium channels (VGSCs) are transmembrane proteins that allow the influx of Na^+^ into excitable cells, including neurons^1,2^. Na_V_1.2, whose alpha subunit is encoded by the gene *SCN2A*, is a VGSC with roles in the generation, propagation, and backpropagation of action potentials in glutamatergic neurons^3–5^. *SCN2A*-related disorders manifest in a highly heterogeneous set of clinical diagnoses. Currently, *SCN2A* is the highest monogenetic risk gene for autism spectrum disorder^4,6,7^, and it has been linked to increased risk for many neurological and neurodevelopmental disorders, including epileptic encephalopathies (EE)^8–11^, intellectual disability (ID)^12,13^, and ataxia^14^. Genetic variation in *SCN2A* has also been linked to bipolar disorder^15–17^ and schizophrenia^18–20^

Mutations in *SCN2A* are generally classified into gain-of-function (GOF) or loss-of-function (LOF) based on their effects on channel expression and biophysical properties. GOF mutations increase sodium influx which promotes increased neuronal excitability and can underlie disorders such as early-onset epilepsy^7,11,21^. On the other hand, LOF mutations reduce the expressed sodium current, whether through reduced trafficking, activity, or protein truncation. LOF variants are often associated with autism spectrum disorder (ASD) and ID, sometimes accompanied by seizures^4,13^. Somatic premature termination codons (PTCs) – non-native stop codons in the protein’s reading frame – are one type of LOF mutation. PTCs, also called “nonsense” mutations, are hypothesized to result in Na_V_1.2 haploinsufficiency, since one copy of the gene is unaffected and the PTC bearing copy generates mRNA presumed to undergo nonsense-mediated decay^22,23^. While this is a commonly held assumption, the effects of different *SCN2A* PTCs on neuronal and behavioral phenotypes remain understudied.

Findings from existing LOF mouse models broadly show that loss of Na_V_1.2 impairs performance in standard learning and memory assays, but results for other behavioral assays are variable, sometimes within the same mouse model^24^. Currently, there are few mouse models with patient-derived *Scn2a* mutations^25–27^, with only one being a nonsense mutation^28^. This highlights the importance of studying individual patient-derived mutations, since not all LOF models show similar phenotypes and the question of whether all truncating variants yield similar outcomes remains understudied, especially since many are rare and *de novo*.

Here, we report the generation and characterization of two novel mouse models carrying the patient-derived PTC variants *SCN2A-*p.Tyr84X (Y84X) and-p.Arg1627X (R1627X). The Y84X variant results in an early truncation in the N-terminal cytoplasmic region, while the R1627X mutation occurs within the fourth transmembrane domain of the protein. We assessed the effects of each mutation on mRNA, Na_V_1.2 protein, and neuronal excitability using electrophysiological techniques, as well as a battery of behavioral analyses. Surprisingly, NMD in the brains of these PTC mouse lines was incomplete (Y84X) or absent (R1627X), suggesting the process of NMD itself may be more highly regulated in brain than previously presumed. Further, both lines showed decreased total Na_V_1.2 protein but at levels statistically above 50%, as would be assumed for pure haploinsufficiency. Slice electrophysiology revealed allele-specific slowing of AP upstroke compared to WT, as well as a threshold difference confined to *Scn2a^Y84X/+^*, consistent with allele-dependent impacts on spike-initiation dynamics.

Our behavioral assessment revealed a mix of distinct phenotypes across both lines, including increased exploration, diminished motor learning, and sex-and genotype-dependent decreased mortality in an induced seizure assay. Our data suggest that PTC mutations in Na_V_1.2 result in cellular and behavioral phenotypes that are both overlapping and mutation-specific, challenging the assumption that all PTCs would lead to similar phenotypes regardless of their genomic location.

## Materials and methods

### Animals

The Y84X and R1627X mutations are found in *SCN2A* patients^29^ (Y84X: https://www.ncbi.nlm.nih.gov/clinvar/variation/VCV000984819.2; R1626X: https://www.ncbi.nlm.nih.gov/clinvar/variation/410985). Due to a single amino acid insertion in the mouse *Scn2a* gene, R1627X in mice is equivalent to R1626X in humans. Transgenic mice were generated in collaboration with the Iowa Genome Editing Core using CRISPR/Cas12 (*Scn2a^Y84X/+^*) and CRISPR/Cas9 (*Scn2a^R^*^1627^*^X/+^*). Both lines were generated on the C57BL/6J strain from Jackson Labs (Strain #000664, Bar Harbor, ME). Transgenic mouse generation and validation details can be found in the Supplementary Material. All experiments were conducted according to the National Institutes of Health guidelines for animal care and were approved by the Institutional Animal Care and Use Committee at the University of Iowa.

## Biochemistry

### Nonsense-Mediated Decay

Whole brains were harvested from adult mice (*n* = 3 per genotype; *Scn2a*^+/+^, *Scn2a^Y84X/+^*, *Scn2a^R1627X/+^*), flash-frozen, and homogenized in TRIzol reagent (Thermo Fisher Scientific). Total RNA was isolated according to the manufacturer’s protocol, followed by lithium-chloride precipitation and ethanol washes. Purified RNA was converted to cDNA using the High-Capacity cDNA Reverse Transcription Kit (Thermo Fisher Scientific). Locus-specific PCR was then performed to amplify (i) exons 1–6 of *Scn2a* from *Scn2a^Y84X/+^* samples (capturing codon 84) and (ii) exon 27 from *Scn2a^R1627X/+^* samples (capturing codon 1627). PCR products were purified and Sanger sequenced by Azenta Life Sciences (South Plainfield, NJ), and amplicon next-generation sequenced by Plasmidsaurus (Louisville, KY) for allele-fraction quantification. Primer sequences and PCR conditions are provided in the Supplementary Material.

### Na_V_1.2 protein expression levels

Mice were euthanized at postnatal day 30 via isoflurane anesthesia followed by decapitation, with full anesthesia confirmed by toe and tail pinch. Brains were rapidly extracted, and one hemisphere was homogenized in RIPA lysis buffer (Thermo Fisher Scientific) supplemented with cOmplete™ Mini EDTA-free protease inhibitor tablets (Roche) to isolate total protein. Protein concentration was quantified using a Bradford assay (Bio-Rad), and 0.026 mg/mL of protein lysate per sample was loaded and analyzed using the Jess™ Simple Western™ system (ProteinSimple, Bio-Techne) according to the manufacturer’s instructions. Detection of Na_V_1.2 was performed using a mouse monoclonal anti-SCN2A antibody (clone 5H10.2, Millipore Sigma). Normalization was completed by Total Protein Assay using Jess™ Simple Western™ system.

### *Ex Vivo* Electrophysiology

On patching days, the order of experimentation (mouse line and genotype) was determined by coin flip to randomize. Acute coronal brain slices (250 µm) containing medial prefrontal cortex were prepared from P25-35 mice. After isoflurane anesthesia and rapid brain removal, slices were cut in ice-cold bath solution and allowed to recover > 30 minutes at room temperature and 5% CO_2_ before recordings at 32 °C.

Neurons were visualized under differential interference contrast (DIC) optics. Recordings were obtained from layer 5b thick-tufted pyramidal-tract (PT) neurons positioned ∼1.5 mm ventral to the dorsal pia and ∼450 µm from the midline in coronal sections. PT neurons were identified physiologically by a rebound depolarization that exceeded resting membrane potential following a strong hyperpolarizing step (−400 pA, 120 ms) with a rebound within 90 ms of step offset^30^.

The external solution contained (in mM): 125 NaCl, 25 glucose, 21.5 NaHCOL, 1 MgClL·6HLO, 2.5 KCl, 1.25 NaHLPOL (anhydrous), and 2 CaClL·2HLO; osmolarity, 302 mOsm. Patch pipettes (Warner Instruments, G150TF-4) were pulled to 3-4 MΩ and filled with (in mM): 113 K-gluconate, 9 HEPES, 4.5 MgClL·6HLO, 10 sucrose, 14 Tris-phosphocreatine, 4 NaL-ATP (monohydrate), 0.3 Tris-GTP, and 0.1 EGTA; pH 7.2–7.5; 285 mOsm.

Whole-cell recordings were made with MultiClamp 700B amplifiers (Molecular Devices) controlled by Clampex and analyzed in Clampfit. Signals were sampled at 50 kHz and low-pass filtered at 20 kHz. Pipette fast capacitance was compensated to 50% under giga-seal conditions before break-in; after establishing whole-cell mode and bridge balance was applied. Series resistance was <12 MΩ for all included recordings; whole-cell capacitance was <12 pF. Action potentials (APs) were evoked with 300-pA, 300-ms depolarizing steps. For biophysical analyses, AP threshold and the maximum rate of rise (peak dV/dt) were quantified from the first spike in a train. Threshold was defined as the membrane potential at which dV/dt first exceeded 13 V sL¹; somatic peak dV/dt was taken as the maximal derivative during the upstroke. The experimenter was blinded to genotype during data analysis.

### General Behavioral Procedures

Mice were housed under regular light cycle with lights on/off at 0900/2100 DST (0800/2000 non-DST). All experiments were conducted during the animals’ light cycle. Mice were same-sex housed in mixed-genotype groups (2–5 mice per cage) on vented cage racks with nesting material as enrichment. Mice were provided with food and water *ad libitum*. Mice were acclimatized for 30-min in the room where the assay was conducted before initiating the test. Equipment was cleaned between trials with 70% ethanol, and the experimenters wore appropriate protective personal equipment when handling the mice. Experimenters and raters remained blinded to genotype throughout behavioral testing and scoring.

### Elevated Zero Maze

Mice were placed in a beige ABS plastic maze (San Diego Instruments, San Diego, CA) elevated 50.0 cm above the table, with an internal diameter of 50.8 cm and an outer diameter of 60.96 cm (internal pathway 5.0 cm wide). The walls on the closed sections were 15.24 cm, and the lip on the open sections was 1.0 cm high. Above the maze, a 45.7 cm ring light (Inkeltech, HaiNing Da Mai E-Commerce Co., China) was positioned, resulting in ∼245 lux in the open sections of the maze. Each mouse underwent a single 5 min trial. Activity was tracked using EthoVision XT 17 software (Noldus, Wageningen, The Netherlands) for distance traveled, velocity, and duration spent in open/closed sections. Mice were excluded from the analysis if they failed to stay on the maze for the entire duration of the trial.

### Open Field Test

Mice were placed in a custom TAP black plastic arena (L × W × HL= 40.0 cm × 42.0 cm × 30.0 cm) over a P95 matte acrylic table. A 45.7 cm ring light was mounted overhead, resulting in ∼115 lux (center) and ∼100 lux (corners). Each mouse underwent a single 10 min trial. Activity was tracked by EthoVision XT 17 software for distance traveled, velocity, and thigmotaxis (the tendency to stay at the edge of the arena). For the latter, the arena was divided into the periphery and the center, each comprising 50% of the total surface area of the arena. Grooming was manually scored by an independent, blind observer from videos with the Behavioral Observation Research Interactive Software (BORIS)^31^.

### Rotarod

Mice were handled for 3 consecutive days before the assay. During the assay, mice were placed on the rotating drum of an accelerating rotarod (Ugo Basile, Varese, Italy). The rotarod accelerated from 4 to 40 rpm over a 5 min period. Mice were given 3 trials/day for 5 consecutive days, with a maximum trial duration of 5 min and an inter-trial interval of at least 10 min. Latency to fall or second passive rotation was recorded for each mouse each day.

### Three-Chamber Social Task

Mice were placed in a matte, black plastic rectangular arena (L × W × HL=L51.0LcmL×L25.4LcmL×L25.4Lcm) divided into three compartments, withLan 11.0Lcm wide opening between compartments and one empty, clear acrylic perforated cylinder in the center of each outer compartment. Mice were first allowed to habituate to the arena for 10 min. Following habituation, mice were placed in a clean cage, while experimenters placed a novel mouse, matched for strain, sex, age, and weight (+/-5.0 g), under one cylinder and a novel, plastic block under the other. Placement of the novel object and novel mice was randomized. After, the experiment mice were allowed to explore the arena for 10 min. Mice were placed in the middle chamber at the beginning of each phase. Activity was tracked with EthoVision XT 17 software to record distance traveled and time spent in each compartment of the arena during both the habituation and sociability phases, as well as the frequency of direct interactions with the cylinders during the sociability phase. Mice were excluded from the sociability phase data analysis if they failed to stay within the arena during the trial.

### Prepulse Inhibition

Mice were restrained in a clear acrylic cylinder (length: 12.7 cm × inner diameter: 3.8 cm) inside a startle response box (L × W × HL=L33.0LcmL×L33.0LcmL×L33.0Lcm. SR-LAB, San Diego Instruments, San Diego, CA). An accelerometer attached underneath the tube restraint recorded movements. Mice were habituated to the isolation cabinet for 5 min with a background white noise of 65LdB, which remained consistent throughout the entire experiment. Each mouse underwent 72 trials in total, in which the first and last 6 trials (Block I and Block IV) consisted solely of pulse-only trials to verify that no habituation to the startle pulse occurred during the experiment. All trials had a randomly spaced inter-trial interval ranging from 8-23 sec. The startle pulse was set to 120LdB, and the prepulse intensities were set to 70 dB (PPI 5), 75 dB (PPI 10), and 80 dB (PPI 15). Each startle tone was 40 msec, and each prepulse was 20 msec. The prepulse preceded the startle by 80 msec, and data were recorded for 65 msec after the startle started. The startle response was recorded in millivolts in SR-LAB software. Percent prepulse inhibition was calculated, normalized to the startle response of the pulse alone trials from Blocks II and III, as follows: percent prepulse inhibitionL=L(startle response for pulse aloneL−Lstartle response for pulse with prepulse)/startle response for pulse alone × 100.

### EEG and EMG Headmount Implantation and Monitoring

Procedures followed published methods^32^. Briefly, mice were anesthetized with isoflurane (3% for induction, 1-1.5% for maintenance) and placed in a stereotaxic frame. Eyes were protected with ophthalmic ointment, meloxicam was given (2 mg/kg, s.c.), and body temperature was maintained on a heating pad. After a midline scalp incision, the skull was cleaned and dried. A modular three-channel head-mount (2 EEG/1 EMG) (8201, Pinnacle Technology, Lawrence, KS) was centered on the skull so that the front holes of the headmount were situated between the frontonasal and coronal sutures and the rear holes of the headmount were between the coronal and lambdoid sutures. Four burr holes were drilled and four stainless steel screws were secured; EMG electrodes were inserted bilaterally into the nuchal muscles. The headmount was secured with dental acrylic, the skin was sutured, and meloxicam was continued for 2 days. Mice recovered for 5 days before recording.

For continuous recordings, mice were placed in a circular mouse cage (8288, Pinnacle Technology). A preamplifier and commutator (Pinnacle Technology) were attached to the headmount. The signals were amplified 100 times, band-pass filtered between 0.5-300 Hz for EEG and 10-300 Hz for EMG and digitized at 1000 Hz via a data acquisition/conditioning system (8206, Pinnacle Technology). Analyses were done with Sirenia Seizure Pro software. Spontaneous seizure activity was analyzed in the EEG2 channel using a 20 sec analysis window with a 1 sec step size. The match was set to > 100% and no upper bound limits were applied. Seizures were detected using the line length method, inputting different thresholds until the highest number of seizure events were marked. All identified seizure events were manually verified.

### Maximal Electroshock (MES) Seizure Induction

This protocol has been described previously^32^. In short, mice were acclimated to the apparatus and saline-moistened ear-clip electrodes for ≥3 h/day on two consecutive days. During testing, current was increased from 0 mA in 1 mA steps every 2 min until a generalized tonic–clonic seizure with hind-limb extension occurred. Outcomes included mortality, seizure-free proportion, current at seizure induction, and extension-to-flexion (E/F) ratio, defined as time in hind-limb extension (>90°) divided by time in flexion (≤90°), scored from video.

### Statistics

Data were graphed and analyzed using OriginPro (OriginLab, Northampton, MA), GraphPad Prism 10.5 (GraphPad Software; San Diego, CA) and RStudio (R4.4.0, afex 1.4-1, emmeans 1.11.1, lme4 1.1-37, lmerTest 3.1-3, rcompanion 2.5.0, survival 3.8-3). Data were analyzed using the statistical tests noted in the results and figure legends (two-way ANOVA, ranked two-way ANOVA, Fisher’s exact test, Chi-square test of independence, log-rank test, linear mixed effects model, or linear model with appropriate follow-up testing). Data are shown graphically as meanL±Lstandard error of the mean (SEM) or standard deviation (SD) for each group. All behavior data were analyzed to assess for sex as a biological variable. Results were considered significant when *p* < 0.05.

## RESULTS

### Assessing NMD in *Scn2a* PTC mouse models

Total RNA was isolated from whole brains of adult *Scn2a* mice, reverse-transcribed, and the premature-termination codon (PTC)–containing regions were amplified for sequence confirmation and quantification (Fig.1A). Amplicons spanning exons 1–6 (capturing codon 84) from *Scn2a^Y84X/+^* and exon 27 (capturing codon 1627) from *Scn2a^R1627X/+^* were PCR-amplified and Sanger-sequenced. At the Y84 locus, chromatograms showed mixed bases at cDNA nucleotide 252 (C/A), yielding TAC (Tyr) in one allele and TAA (PTC) in the other (Fig.1B). At the R1627 locus, chromatograms showed mixed bases at cDNA nucleotide 4879 (C/T; A/G on the reverse strand), corresponding to CGA (Arg) and TGA (PTC), respectively. These data confirm the presence of the expected heterozygous PTCs in both models (Fig.1B).

**Figure 1.**
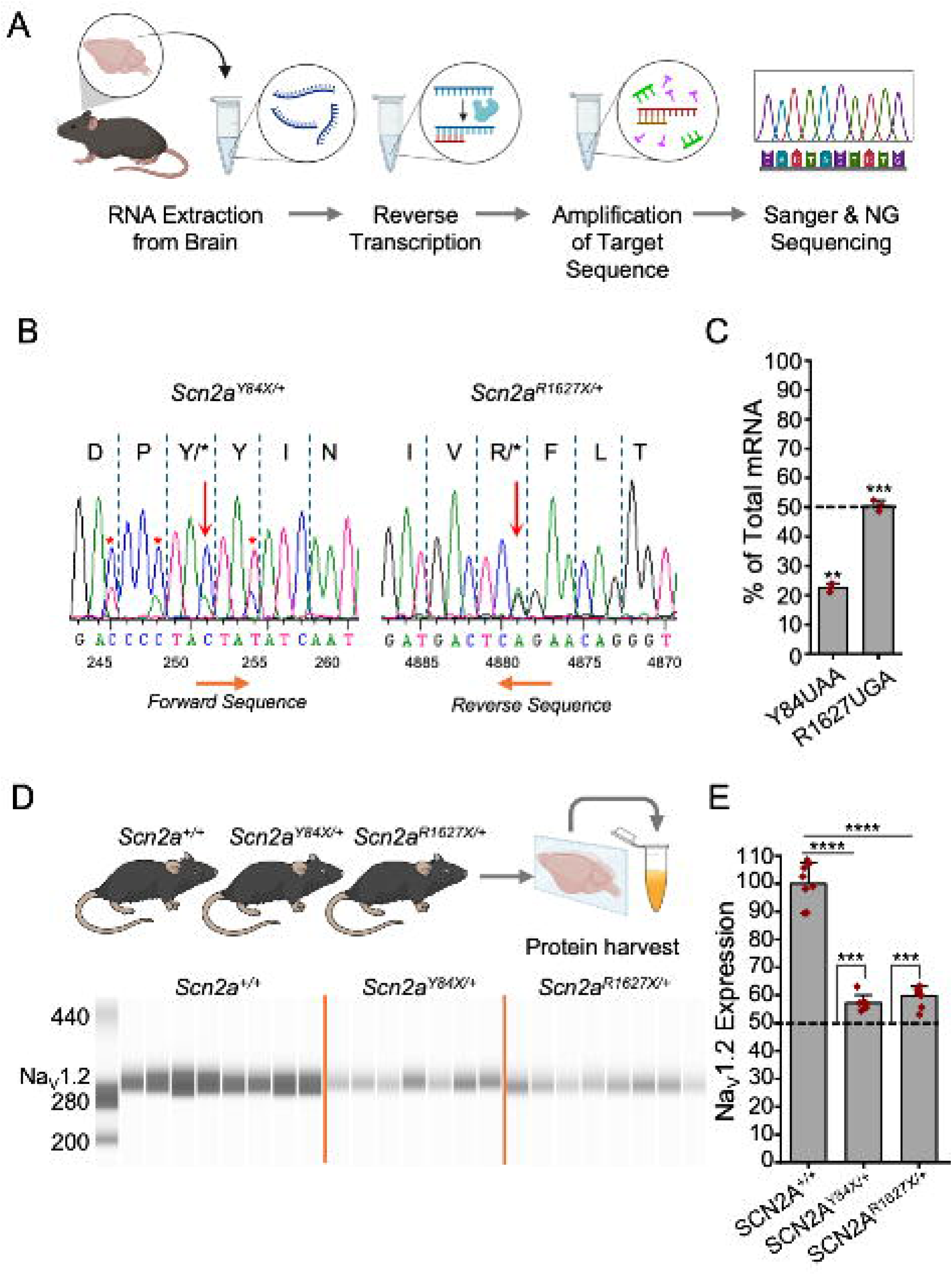
Biochemical characterization of *Scn2a* mouse models. **(A)** Schematic representation of experimental design performed to quantify NMD in *Scn2a* mouse models. **(B)** Sanger sequencing of *Scn2a* mRNA shows the Y84UAA mutation at 252bp (Left) and R1626UGA at 4879bp (Right). **(C)** NGS quantifies transcripts containing PTCs in *Scn2a*^Y84X/+^ and *Scn2a*^R1627X/+^ (n = 3 mice per genotype, *p* < 0.001). **(D)** Representative Jess automated Western blot showing expression of Na_V_1.2 in half-brain lysates. **(E)** Bar graph quantification of band intensity shown in **(D)** displaying decreased Na_V_1.2 protein in mutant brains compared to WT littermates (ANOVA, *p* < 0.0001) but over 50% (One-sample t-test, *p* < 0.001). Data are mean ± SD (n = 7-8 mice per genotype). SCN2a antibody used Millipore Sigma clone 5H10.2.

Allele fractions were then quantified by targeted next-generation sequencing (NGS) per mouse and averaged across animals (Fig.1C). The mutant-allele fraction averaged 22.5 ± 0.8% in *Scn2a^Y84X/+^* brains and 50.49 ± 1.0% in *Scn2a^R1627X/+^*brains (*p* < 0.001), confirming expression of the PTC-bearing transcripts in both genotypes. Relative to the predicted allelic balance (50%), the Y84X mutant allele was under-represented (22.5%, *p* < 0.001), consistent with incomplete NMD at this site. By contrast, R1627X-bearing transcripts were at balance (50.49%, *p* > 0.6), suggesting no detectable NMD occurred to this PTC-bearing transcript.

### Quantification of Na_V_1.2 in heterozygous *Scn2a* PTC mice

Na_V_1.2 protein abundance was quantified in half-brain lysates from P30 *Scn2a* mouse models using the Jess automated Western blot platform, with total-protein normalization (Jess total-protein assay) and normalization to the wild-type (WT) mean (Fig. 1D). Across WT (*n* = 8), *Scn2a^Y84X/+^* (*n* = 7), and *Scn2a^R1627X/+^* (*n* = 8) cohorts, Na_V_1.2 levels differed significantly (*p* < 0.0001). Relative to WT, Na_V_1.2 was reduced to 57.2 ± 2.84% in *Scn2a^Y84X/+^* and 59.6 ± 3.51% in *Scn2a^R1627X/+^* (mean ± SD) (Fig. 1E). Despite this reduction, Na_V_1.2 remained > 50% of WT in each mutant line (*p* < 0.001 for both), with no significant difference detected between the two variants.

### Electrophysiological assessment of *Scn2a* PTC mouse models

Since Y84X and R1627X variants introduce PTCs in the *Scn2a* gene, P25-P30 *Scn2a* mouse acute mPFC slices were examined by performing whole-cell current clamp on layer 5b pyramidal tract (PT) neurons. To ensure recordings were made from PT neurons, we used well-established electrophysiological criteria. In acute mPFC slices, corticofugal layer 5b neurons can be distinguished by their high expression of HCN channels that manifest as voltage ‘sag’ visible during a strong hyperpolarization step, and a rebound that peaks to a more depolarized potential than V_m_ at rest following current offset (see Methods)^5,30,33^.

Action potentials (APs) were evoked with a 300 pA, 300 ms depolarizing step (Fig. 2A). Phase-plane plots of dV/dt versus membrane voltage were used to quantify the kinetics of the rising phase (Fig. 2B). This approach differentiates contributions of AIS and somatodendritic sodium channels^5^.

**Figure 2.**
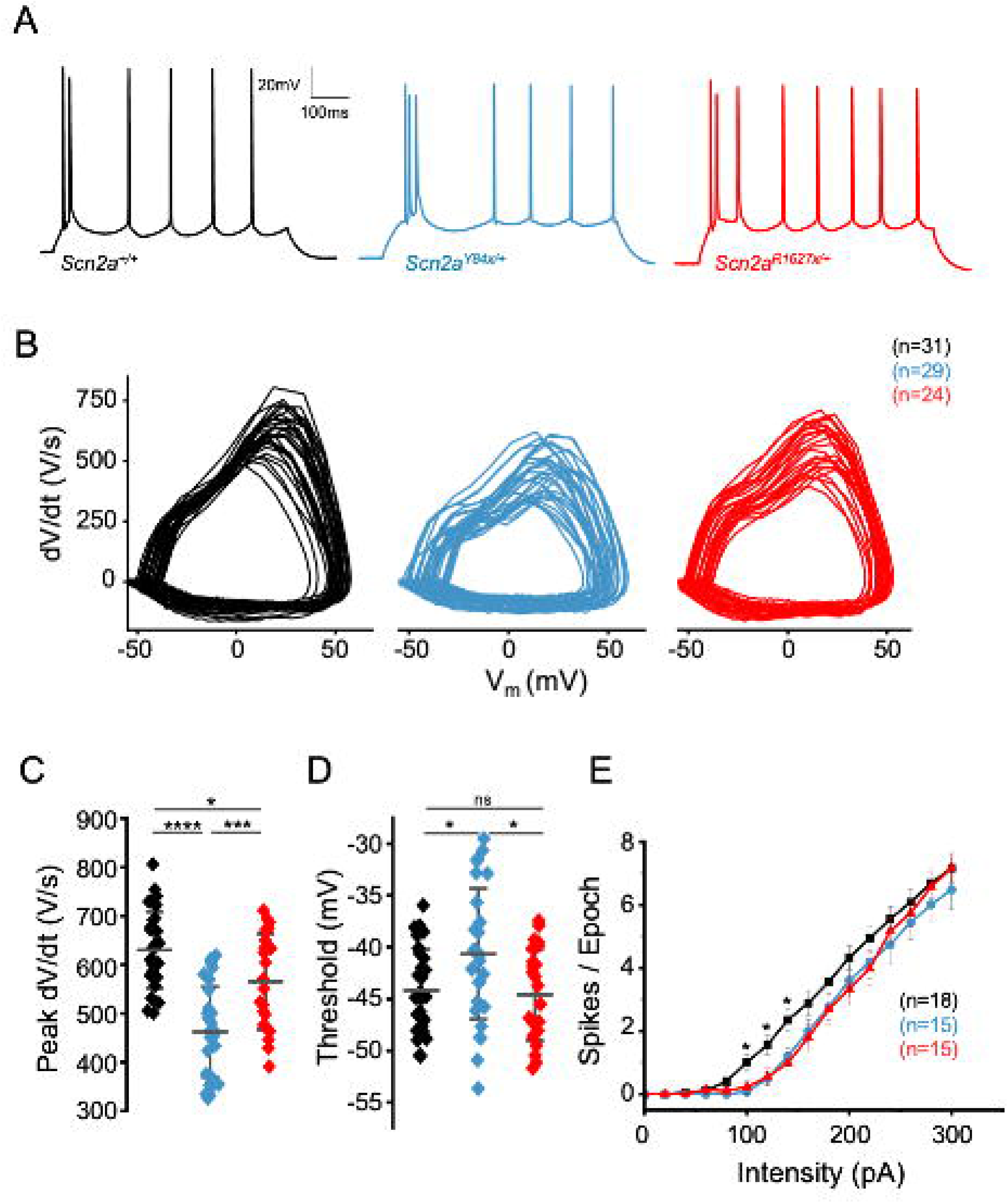
Electrophysiological characterization of *Scn2a* mouse models. **(A)** Action potentials generated by current injection (300pA, 300ms) in *Scn2a*^+/+^ (black), *Scn2a*^Y84X/+^ (cyan) and *Scn2a*^R1627X/+^ (red) cortical pyramidal neurons. **(B)** Phase-plane plots of all action potentials recorded (same color-coding as **(A)**). **(C)** Maximum speed of the rising phase of the APs. **(D)** Threshold of APs (Data are mean ± SD. One-way ANOVA, followed by Tukey’s post hoc test for multiple comparisons. **(E)** F–I curves of cortical pyramidal neurons from WT (*n* = 18), *Scn2a*^Y84X/+^ (*n* = 15), and *Scn2a*^R1627X/+^ (*n* = 15) mice showing mean spikes ± SEM. Group differences were assessed per step by one-way ANOVA with Tukey’s HSD post hoc tests.

Somatic peak dV/dt differed markedly across genotypes. Neurons from WT mice exhibited a rapid upstroke, with a mean of 630.3 ± 78.47 V sL¹ (mean ± SD, *n* = 31 cells) (Fig. 2C), while neurons expressing *Scn2a^R1627X/+^* exhibited slower kinetics (564.9 ± 98.3 V sL¹, *n* = 24; *p* < 0.05). On the other hand, APs recorded from *Scn2a^Y84X/+^* neurons were significantly slower than both WT (*p* < 0.0001) and *Scn2a^R1627X/+^* (*p* < 0.001) (462.1 ± 92.7 V sL¹, *n* = 29). Although these reductions are consistent with previous observations that Na_V_1.2 heterozygous LoF mutations attenuate the velocity of membrane depolarization, the more pronounced reduction in *Scn2a^Y84X/+^* neurons suggests that the Y84X mutation may result in a more severe functional deficit than R1627X^34,35^.

AP spike threshold, measured from phase-plane plots at the first rapid rise in dV/dt, was also assessed. Thresholds were similar in wild-type neurons (−44.2 ± 3.9 mV) and *Scn2a^R1627X/+^* neurons (−44.5 ± 4.4 mV; *p* > 0.05) (Fig. 2D). In contrast, *Scn2a^Y84X/+^* neurons required slightly greater depolarization to spike (−40.7 ± 6.3 mV, *p* < 0.05 vs wild type and *Scn2a^R1627X/+^*), indicating that this PTC allele shifts the AP threshold to a more depolarized potential (Fig. 2D).

To evaluate how PTC alleles affect neuronal excitability, we injected incrementing 300 ms current steps (0–300 pA) and counted the number of spikes per step (F–I curves, Fig. 2E). Both PTC models fired fewer spikes than wild-type at near-rheobase inputs (100–140 pA). At higher current injections however, firing in *Scn2a* PTC neurons converged toward the wild-type phenotype.

### Weight

We explored baseline health characteristics of heterozygous *Scn2a* PTC mice. These mice were viable as heterozygotes and inheritance of the *Scn2a* PTC variants followed expected Mendelian ratios. Prior to behavioral testing, we observed an effect of genotype on weight across both lines, such that *Scn2a* PTC mice were slightly smaller than their respective WT littermates (Y84X *n* = 37, *p* < 0.01; R1627X *n* = 33, *p* < 0.01). *Scn2a^Y84X/+^* mice were on average 2.03g smaller than WT (Fig. S1A), while *Scn2a^R1627X/+^*mice were 2.59g smaller on average (Fig. S1B). As expected, we also observed a main effect of sex on weight in both lines, with female mice being smaller than males on average. In the Y84X line, females were 6.81g (± 0.489, *n* = 36, *p* < 0.01) smaller than males, while in the R1627X line, females were 7.06g (± 0.488, *n* = 37, *p* < 0.01) smaller than males (Supp Table 1). Overall, we would not expect these weight differences to impede the animals’ ability to perform behavioral tasks.

### Locomotor and exploratory behavior

We examined whether *Scn2a^Y84X/+^* and *Scn2a^R1627X/+^*mice display abnormal locomotor and exploratory behaviors. In both the elevated zero maze and open field test, we observed no differences in distance traveled by *Scn2a^Y84X/+^* (Fig. 3A, E) and *Scn2a^R1627X/+^* mice compared to their corresponding WT littermates (Fig. 3C, G). We observed a main effect of sex on distance traveled in the elevated zero maze in the R1627X line, such that males traveled greater distances than females (Supp Table 2). However, this was not observed in the open field test, nor the Y84X line in either the elevated zero maze or the open field test (Supp Table 2). These data suggest that Na_V_1.2 deficiency does not cause major impairments in baseline locomotion or exploration.

**Figure 3.**
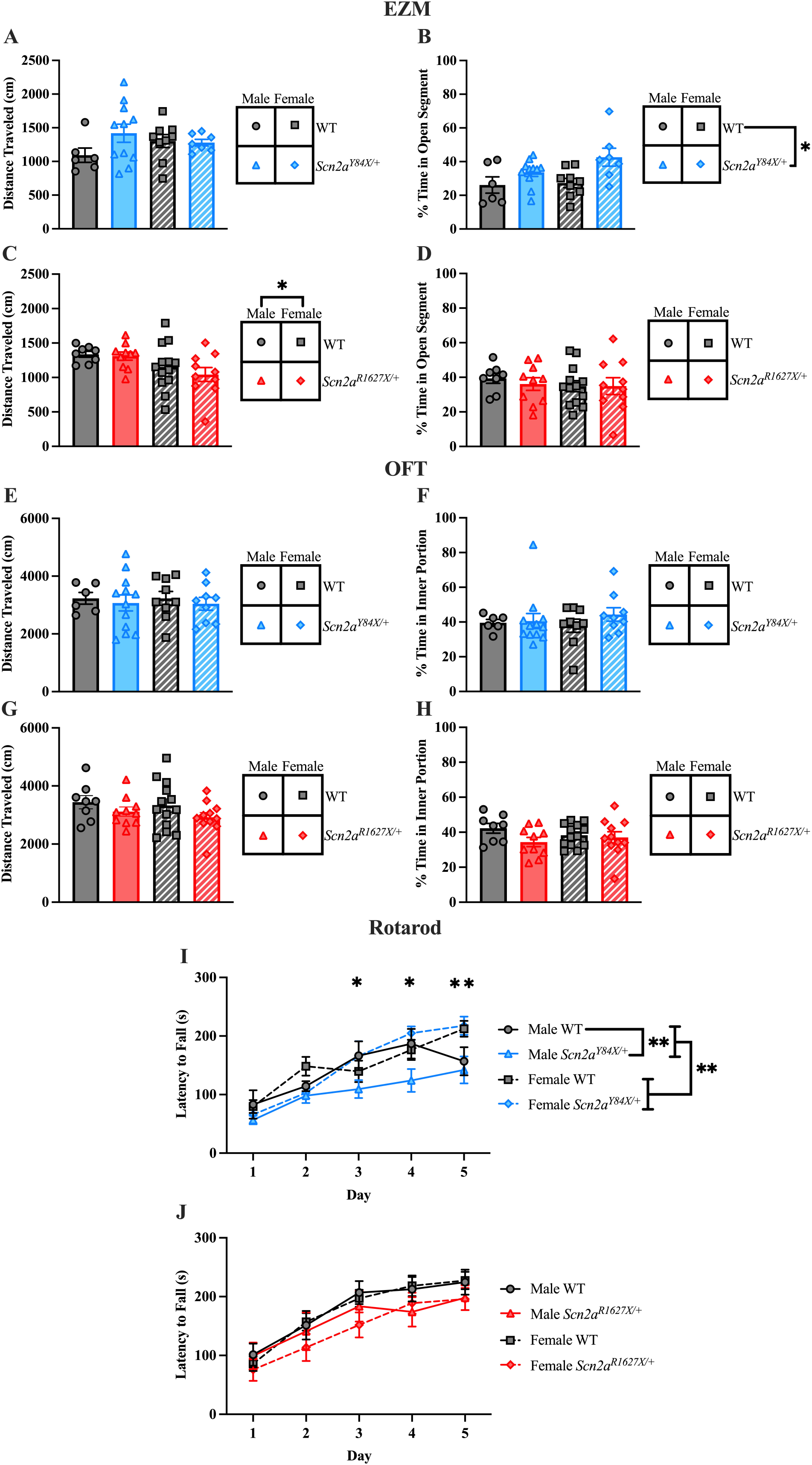
*Scn2a^Y84X/+^*mice exhibit diminished anxiety-like behaviors and alterations in motor behavior. (**A)** There were no differences in distance traveled in the Y84X line for the elevated zero maze. **(B)** *Scn2a^Y84X/+^*mice spent significantly more time in the open segment of the elevated zero maze compared to WT mice. **(C)** Male mice in the R1627X line traveled longer distances in the elevated zero maze. **(D)** There were no differences in time spent in the open segment of the elevated zero maze in the R1627X line. **(E)** There were no differences in distance traveled or **(F)** time spent in the inner portion of the arena of the open field task in the Y84X line. **(G)** There were no differences observed in distance traveled or **(H)** time spent in the inner portion of the arena of the open field task in the R1627X line. **(I)** We observed a genotype × session × sex interaction in the rotarod assay for the Y84X line. Post-hoc testing showed a difference between male *Scn2a^Y84X/+^* and WT mice on sessions 3 and 4. Further, we observed a sex × session interaction effect between males and females on session 5, as well as a main effect of session. **(J)** No differences in rotarod performance were detected between mice in the R1627X line. All genotypes improved over the course of training. **(A-J)** Data are expressed as mean ± SEM. **(A-B)** *n* = 33 (WTs: *n* = 6 males, *n* = 9 females; *Scn2a^Y84X/+^*: *n* = 11 males, *n* = 7 females). **(C-D)** *n* = 41 (WTs: *n* = 8 males, *n* = 13 females; *Scn2a^R1627X/+^*: *n* = 10 males, *n* = 10 females). **(E, F, I)** n = 36 (WTs: *n* = 6 males, *n* = 9 females; *Scn2a^Y84X/+^*: *n* = 12 males, *n* = 9 females). **(G, H, J)** *n* = 42 (WTs: *n* = 8 males, *n* = 13 females; *Scn2a^R1627X/+^*: *n* = 10 males, *n* = 11 females).

### Motor performance learning

We hypothesized that Na_V_1.2 may be involved in motor learning due to its presence in cerebellar granule neurons. Therefore, we tested the ability of mice to learn the accelerated rotarod task. In the Y84X line, we observed a genotype × session × sex interaction, a sex × session interaction, and a main effect of session, with no main effect of sex or genotype (Fig. 3I). Follow-up testing showed no differences in motor learning between female *Scn2a^Y84X/+^* and WT mice, but there was a significant difference between male *Scn2a^Y84X/+^* and WT mice on sessions 3 and 4. Furthermore, follow-up testing showed a significant difference between males and females on session 5. In the R1627X line, we found a main effect of session, but no main effect of sex or genotype, and no interaction effect (Fig. 3J and Supp Table 3). Therefore, the Y84X mutation is associated with a male-predominant impairment in motor performance learning on the accelerating rotarod task, while the R1627X mutation does not seem to impact performance on the task.

### Anxiety-like and grooming behavior

We explored anxiety-like behaviors using the elevated zero maze and the open field test, which measure a mouse’s willingness to explore more exposed parts of their environment. We hypothesized that mutant mice in both lines would spend more time in high-risk areas, due to sensory-seeking behaviors observed in some patients with autism. In the Y84X line, *Scn2a^Y84X/+^* mice spent significantly more time exploring the open portions of the elevated zero maze compared to their WT littermates (Fig. 3B). We observed no main effect of sex or interaction effects. However, we observed two *Scn2a^Y84X/+^* mice jump from the apparatus before completing the trial, and one *Scn2a^Y84X/+^* mouse jumped directly onto the experimenter’s chest before being placed on the assay. These behaviors were not observed in WT littermates from the Y84X line. We observed no genotype or sex effects for time spent in the center of the open field arena (Fig. 3F). In the R1627X line, we observed no differences between groups in the time spent in the open portion of the elevated zero maze (Fig. 3D). However, one *Scn2a^R1627X/+^* mouse dove off the apparatus before completing the 5-min trial, which was not observed in WT littermates from this line. In the open field test, we observed no group differences in the R1627X line in time spent in the center of the arena (Fig. 3H and Supp Table 4). We conclude that the Y84X mutation is associated with lower anxiety-like behaviors in contexts that present higher risk, such as the elevated zero maze, while the R1627X mutation does not have an impact on these behaviors.

We next measured grooming behavior during the open field task. Grooming is an innate behavior, and we hypothesized that *Scn2a* PTC mice would groom more frequently or for longer periods, representative of repetitive and restrictive behavior, a cardinal feature of autism^36^. In the Y84X line, *Scn2a^Y84X/+^* mice showed a nonsignificant trend toward increased grooming frequency, and they spent significantly more time grooming than their WT littermates (Fig. 4A-B). This suggests that once *Scn2a^Y84X/+^* mice initiated a grooming bout, they remained engaged in this behavior for longer periods. In the R1627X line, we observed a genotype × sex interaction in grooming frequency (Fig. 4C). Post-hoc analysis showed no difference between females; however, male *Scn2a^R1627X/+^* mice groomed more frequently than their WT littermates. *Scn2a^R1627X/+^* mice also spent more time grooming than WT littermates (Fig. 4D and Supp Table 5). Together, these findings suggest that both *Scn2a^Y84X/+^* and *Scn2a^R1627X/+^* mice exhibit restrictive and repetitive grooming behaviors.

**Figure 4.**
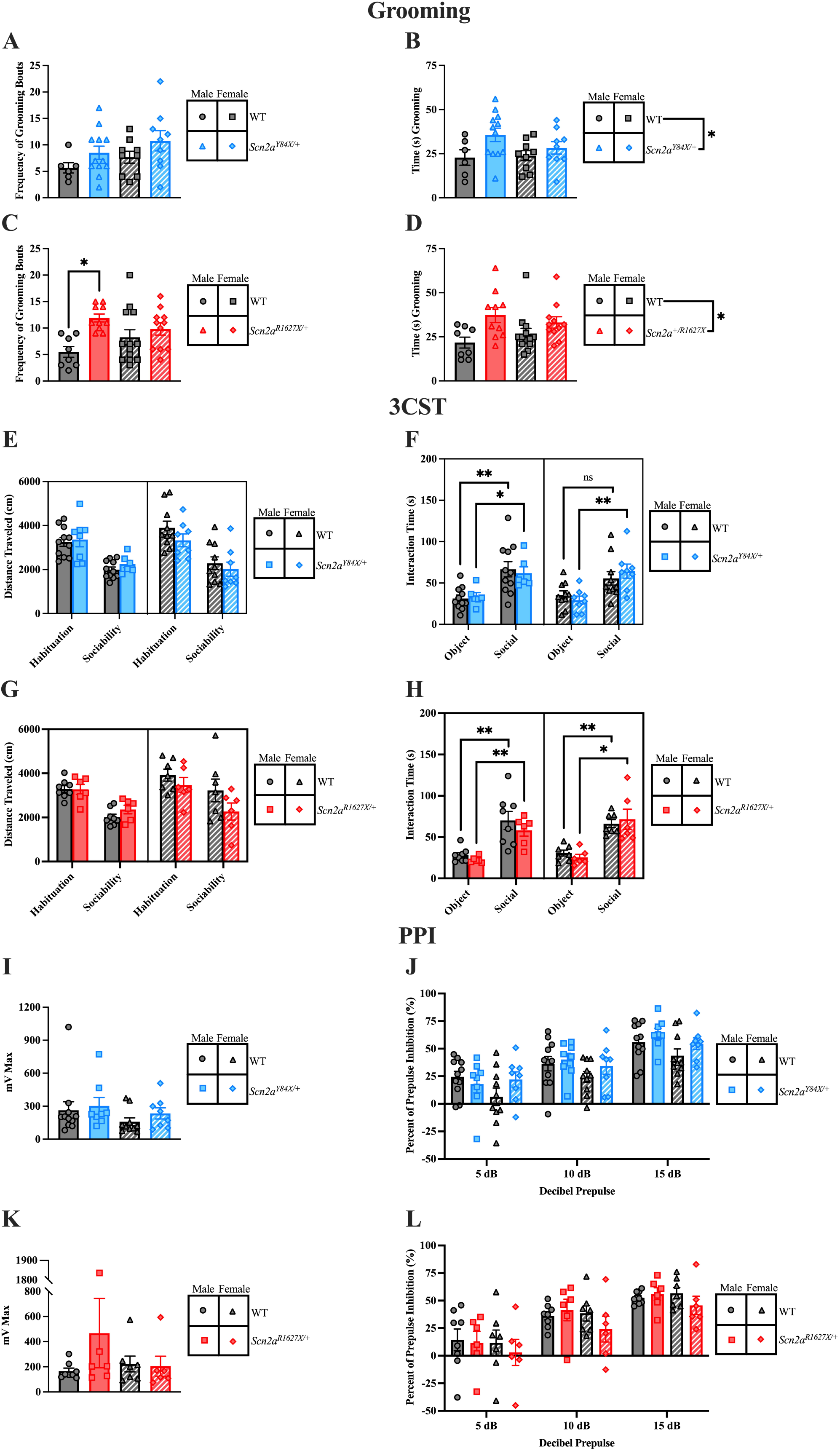
*Scn2a^Y84X/+^* and *Scn2a^R1627X/+^* mice exhibit restrictive and repetitive behaviors. (A) There were no differences in grooming frequency in the Y84X line for the open field test. (B) *Scn2a^Y84X/+^* mice spent significantly more time grooming compared to WT mice. **(C)** *Scn2a^R1627X/+^* male mice had a higher number of grooming bouts compared to WT males. **(D)** *Scn2a^R1627X/+^* mice spent more time grooming in the open field assay. **(E)** There were no differences in distance traveled in the three-chamber social task. **(F)** WT females in the Y84X line did not show preference for the novel mouse in the sociability phase of the three-chamber social task. **(G)** There were no differences observed in distance traveled or **(H)** time spent with the conspecific mice in the R1627X line in the three-chamber social task. **(I)** There were no differences in startle response or **(J)** percent of prepulse inhibition in the Y84X line. **(K)** There were no differences in startle response or **(L)** percent of prepulse inhibition in the R1627X line. **(A-L)** Data are expressed as mean ± SEM. **(A-B)** *n* = 36 (WTs: *n* = 6 males, *n* = 9 females; *Scn2a^Y84X/+^*: *n* = 12 males, *n* = 9 females). **(C-D)** *n* = 42 (WTs: *n* = 8 males, *n* = 13 females; *Scn2a^R1627X/+^*: *n* = 10 males, *n* = 11 females). **(E-F)** *n* = 35 & 37 (WTs: *n* = 11 males, *n* = 10 females; *Scn2a^Y84X/+^*: *n* = 6 & 8 males, *n* = 8 females). **(G-H, K-L)** *n* = 27 (WTs: *n* = 8 males, *n* = 7 females; *Scn2a^R1627X/+^*: *n* = 6 males, *n* = 6 females). **(I-J)** *n* = 37 (WTs: *n* = 11 males, *n* = 10 females; *Scn2a^Y84X/+^*: *n* = 8 males, *n* = 8 females).

### Social preference

Given that mutations in Na_V_1.2 have been linked to autism, we sought to investigate whether PTC mutations in *Scn2a* would affect social preference with the three-chamber social preference task. To control for motor behavior and exploration, we examined the distance traveled during both the habituation and sociability phases of the task. In both mouse lines we observed a main effect of phase, such that all groups were more active during the habituation phase than the sociability phase. However, we found no significant differences between sexes or genotypes in total exploration (Fig. 4E, G and Supp Table 6A). During the social phase in the Y84X line, we observed that, on average, all mice spent more time with the novel mouse than with the novel object; however, there were no effects of genotype or sex (Supp Table 6A). Further, we found that WT females do not exhibit a statistically significant preference for conspecific mice over the object; however, all other groups showed a clear social preference (Fig. 4F and Supp Table 6B). Interestingly, we observed two male *Scn2a^Y84X/+^* mice exit the arenas during the sociability phase. This was not noted in WT littermates or any mice in the R1627X line. In the R1627X line, we observed no differences in the interaction time with each cylinder by genotype, nor were there any interaction effects (Supp Table 6A). All groups spent more time with the novel, conspecific mouse over the novel object (Fig. 4H and Supp Table 6B). Taken together, these data suggest that *Scn2a^Y84X/+^* and *Scn2a^R1627X/+^* mice display social preference similar to WT mice.

### Sensorimotor gating

Given that Na_V_1.2 deficiency has been linked to developmental disorders with impaired sensorimotor gating^37^, we tested startle response and sensorimotor gating using the prepulse inhibition task. *Scn2a^Y84X/+^* and *Scn2a^R1627X/+^* showed normal auditory startle responses compared to their respective WT littermates (Fig. 4I, K), with detectable responses throughout the task. In the Y84X line, all groups exhibited a mild but significant habituation to the startle stimulus between blocks I and IV, whereas the R1627X line showed a similar but non-significant trend (Fig. S2A-B). Further, all groups displayed normal sensorimotor gating, with no significant effects of genotype or sex (Fig. 4J, L, and Supp Table 7). Higher decibel prepulses yielded a higher percent of prepulse inhibition across all mice. We conclude that *Scn2a^Y84X/+^* and *Scn2a^R1627X/+^* mice have adequate startle response and normal sensorimotor gating.

### Spontaneous Seizures and MES Threshold

Continuous EEGs were recorded in *Scn2a* PTC mice to evaluate whether they were prone to spontaneous seizures. We observed rare epileptiform activity in both WT and *Scn2a* PTC mice, with no obvious differences in frequency between genotypes in either mouse line, suggesting that these *Scn2a* PTC mice do not display spontaneous seizure activity to a greater extent than WT littermates. To determine whether stimulation of Na_V_1.2-deficient neurons would increase mortality following seizure induction, we performed a maximal electroshock threshold assay. In the Y84X line, we observed a main effect of genotype for mortality, such that fewer *Scn2a^Y84X/+^* mice died from the MES threshold assay compared to WT mice (Fig. 5A). We did not observe a main effect of sex. The difference in percent mortality is not attributable to the current used, as there was no difference between groups for the current at which a seizure was induced or in the likelihood of remaining seizure-free (Fig. 5C and Fig. S3A). Further, we analyzed extension-to-flexion (E/F) ratio, which is an indication of seizure severity^38^. No differences were noted in the E/F ratio of *Scn2a^Y84X/+^* mice compared to WT (Fig. 5D). In the R1627X line, we also observed that *Scn2a^R1627X/+^* mice were less likely to die from the MES threshold assay than their WT littermates, but there were no effects of sex (Fig. 5B). Furthermore, there were no significant group differences in the current needed to induce a seizure or the possibility of being seizure-free (Fig. 5E and Fig. S3B). Finally, we observed no differences in seizure severity, as analyzed from the E/F ratio (Fig. 5F and Supp Table 8). These results suggest that both the Y84X and the R1627X mutations are protective against seizure fatality in the C57BL/6J genetic background, but do not affect seizure threshold or severity.

**Figure 5.**
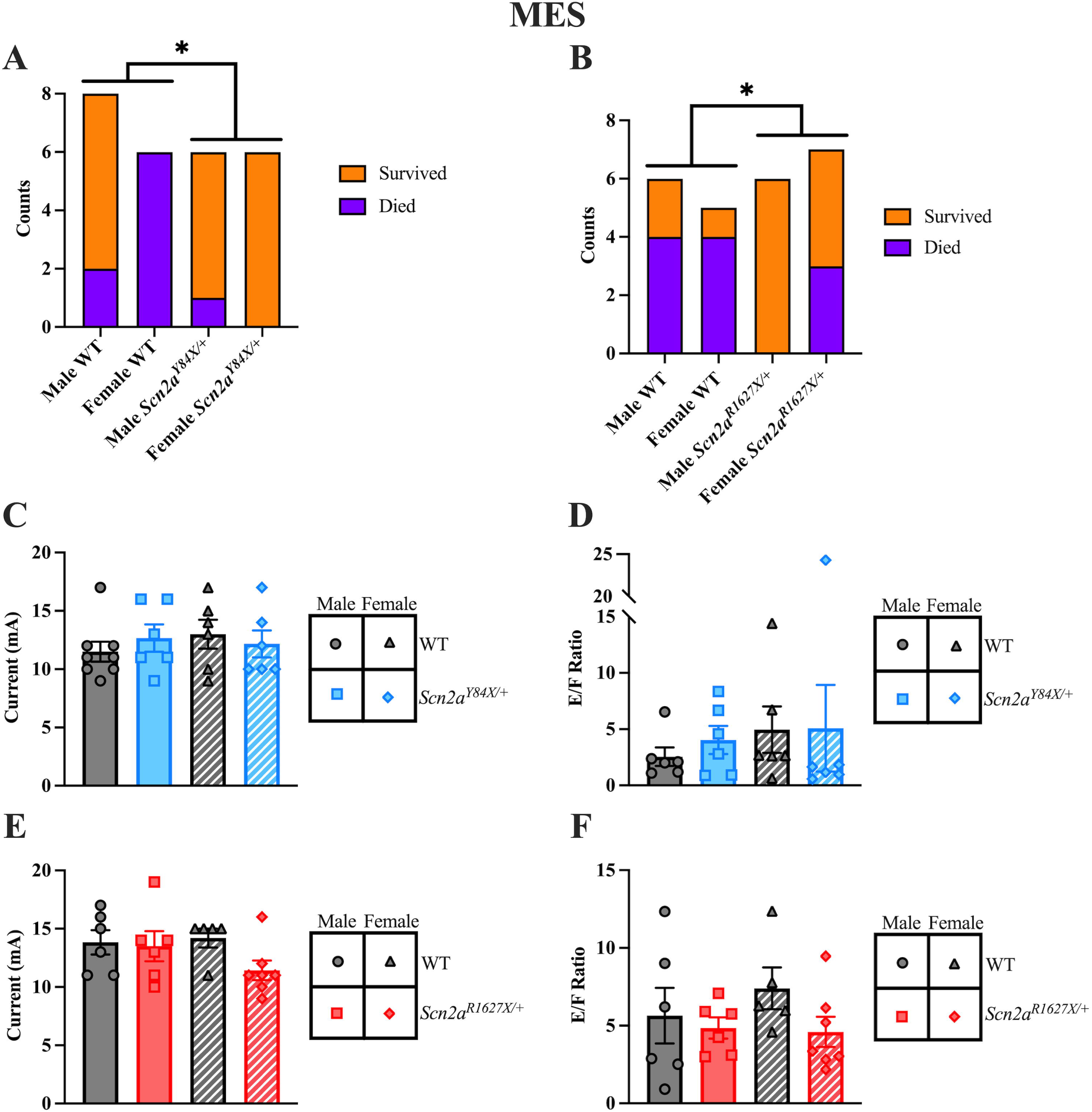
*Scn2a^Y84X/+^* mice show decreased mortality in seizure induction assay. (**A)** No female *Scn2a^Y84X/+^* mice died from the maximal electroshock threshold assay, while all their WT littermates did. **(B)** *Scn2a^R1627X/+^* mice were less likely to die from the MES threshold assay than their WT littermates. **(C)** In the Y84X line, there is no difference in the current at which seizures were induced or **(D)** the severity of the seizures. **(E)** There were no differences in current at which seizures were induced or **(F**) seizure severity in the R1627X line. **(A-F)** Data are expressed as mean ± SEM. **(A, C)** *n* = 26 (WTs: *n* = 8 males, *n* = 6 females; *Scn2a^Y84X/+^*: *n* = 6 males, *n* = 6 females). **(B, E-F)** *n* = 24 (WTs: *n* = 6 males, *n* = 5 females; *Scn2a^R1627X/+^*: *n* = 6 males, *n* = 7 females). **(D)** *n* = 24 (WTs: *n* = 6 males, *n* = 6 females; *Scn2a^Y84X/+^*: *n* = 6 males, *n* = 6 females).

## Discussion

Na_V_1.2 supports important roles in neuronal function and has been linked to multiple neurological and psychiatric disorders. In this study, we characterized biochemical, cellular, and behavioral phenotypes associated with two patient-derived PTC variants in *SCN2A* with two newly generated mouse models: *Scn2a^Y84X/+^* and *Scn2a^R1627X/+^*. We demonstrate that the location of the PTC governs molecular handling and produces allele-specific physiological and behavioral outcomes.

At the mRNA level, *Scn2a^Y84X/+^*-expressing neurons exhibited partial NMD (mutant-allele fraction = 22.5%), a notable finding indicating that an early coding-sequence PTC in *Scn2a* is not fully degraded in the brain (Fig. 1C). In contrast, R1627X transcripts were at allelic balance, consistent with a C-terminal-exon PTC that typically escapes exon-junction– complex–dependent NMD^39^. Despite these RNA differences, Na_V_1.2 protein was similarly reduced in both lines (∼57–60% of WT) (Fig. 1E). This convergence suggests effective haploinsufficiency at the proteome level. Importantly, a PTC allele does not necessarily recapitulate the quantitative protein reduction observed with a clean allele deletion. We observed Na_V_1.2 protein significantly above 50% of WT (*p* < 0.001 for both) rather than a strict 1:1 loss. This difference may influence cellular and behavioral phenotypes and could stem from variable and site-specific read-through at each PTC or compensatory regulation of the WT allele.

Previous work has shown that *Scn2a* haploinsufficiency disrupts dendritic excitability and synaptic function in layer 5b pyramidal tract neurons^5,34^, which makes them an ideal model for dissecting the consequences of *Scn2a* premature stop codons via current-clamp recordings. Physiologically, PTCs demonstrate unique phenotypes. Both PTCs reduce the AP upstroke, but the decrement in peak dV/dt is larger in Y84X than in R1627X-expressing neurons (26.7% vs 10.3% respectively) (Fig. 2C). Moreover, spike threshold is depolarized only in *Scn2a^Y84X/+^*mice, whereas thresholds from *Scn2a^R1627X/+^* neurons remain indistinguishable from WT (Fig. 2D). Consistent with reduced Na_V_1.2 drive near rheobase, both alleles show decreased firing at low input currents (Fig. 2E). However, in this framework, the Y84X variant exerts a greater impact on somatic upstroke kinetics and rheobase-proximal firing. These allele-dependent signatures reinforce that PTC context — not merely the presence of a PTC — governs how Na_V_1.2 insufficiency manifests at the soma/proximal initial segment.

Collectively, these findings show that the position of a PTC in *Scn2a* shapes neuronal physiology. *Scn2a^Y84X/+^* undergoes partial NMD, whereas *Scn2a^R1627X/+^* escapes it, yet both lines exhibit a similar—though not strictly halved—reduction in Na_V_1.2 protein. Functionally, both mutations slow the action-potential upstroke, with a larger decrement and a selective threshold depolarization in *Scn2a^Y84X/+^*. These results indicate that distinct PTC variants are not interchangeable across molecular and cellular levels.

### Behavioral phenotypes associated with *Scn2a* PTCs

Both *Scn2a^Y84X/+^* and *Scn2a^R1627X/+^* mice showed subtle reductions in body weight compared to their respective WT controls. Gross motor activity was preserved across genotypes, and neither line showed impairments in baseline locomotion, similar to a few other haploinsufficient *Scn2a* models, although hyperactivity has also been reported^40–42^. In contrast, *Scn2a^Y84X/+^* mice showed a male-specific impairment in motor learning on the rotarod task, a phenotype reported in one other model^42^ but not observed in *Scn2a^R1627X/+^*mice or most other haploinsufficient models^5,40,41^. This variant and sex-specific impairment may reflect subtle differences in cerebellar function, due to Na_V_1.2’s expression in cerebellar granule neurons^43^.

Additionally, *Scn2a^Y84X/+^* mice spent significantly more time in the open arms of the elevated zero maze, suggesting increased risk-taking behavior, while *Scn2a^R1627X/+^* mice showed no alterations. Further, both *Scn2a^Y84X/+^* and *Scn2a^R1627X/+^*mice had unusual exploratory behaviors, such as jumping or diving from elevated zero maze, or male *Scn2a^Y84X/+^* mice jumping out of the arena in the three-chamber social task. Instances of increased exploratory behavior have been observed elsewhere^5,41,42^, although primarily in the elevated zero or elevated plus maze rather than the open field test, where, similar to our findings, most models show no differences^5,40,41^. These phenotypes could reflect sensory-seeking behaviors, although more careful experimentation focused on sensory function and risk-taking behaviors are required to clarify this.

A key finding was the increase in grooming behaviors in both *Scn2a^Y84X/+^*and *Scn2a^R1627X/+^* mice, suggesting heightened repetitive and restrictive behavior – a core diagnostic feature of ASD. *Scn2a^Y84X/+^* mice spent more time grooming, and while their number of grooming bouts were only trending towards an increase; the long time spent in a single grooming bout suggests behavioral perseveration. *Scn2a^R1627X/+^* mice also had longer grooming duration, and *Scn2a^R1627X/+^* males showed higher grooming frequency than WT controls. Increased grooming behavior has been reported in some juvenile haploinsufficient models, but not in adult mice, except the gene-trap mouse model^5,40,41,44,45^. Our findings suggest that Na_V_1.2 deficiency increases repetitive behaviors into adulthood.

Other domains—sociability in the three-chamber task and sensorimotor gating—were intact in both lines, suggesting a selective rather than global alteration of behavior.

Finally, in the MES assay, both *Scn2a^Y84X/+^* and *Scn2a^R1627X/+^* mice showed reduced seizure-induced mortality, despite having similar seizure thresholds and severity compared to WT littermates. While this superficially suggests that *Scn2a* PTC variants confer protection against fatal seizures, children with *SCN2A* LOF mutations often develop seizure disorders, which can be difficult to treat or fatal. The genetic strain used for both of our *Scn2a* PTC lines, C57BL/6J, is associated with less severe seizure phenotypes in many models of SCN2A-related disorders^4,24–26,28,46,47^, so this particular result should be interpreted with caution.

Together, the behavioral data demonstrate that the Y84X and R1627X variants result in overlapping but allele-specific phenotypes. *Scn2a^Y84X/+^*mice generally show more pronounced changes—mirrored by stronger alterations in slice physiology—in motor and exploratory behaviors. Both *Scn2a^Y84X/+^*and *Scn2a^R1627X/+^* mice display restrictive and repetitive grooming and appear protected against MES-induced mortality. However, only *Scn2a^Y84X/+^* mice display increased exploration and locomotor learning deficits. Overall, these findings demonstrate that distinct PTC variants are not interchangeable, producing partially shared but allele-specific functional outcomes across molecular, cellular, and behavioral levels.

### Challenges, limitations, and future directions

RNA and protein measurements were obtained from whole brain, precluding cell type-specific analyses. NMD and Na_V_1.2 levels may differ in PT neurons versus other neuronal types; cell type-specific measurements would help resolve allele-specific mechanisms. We did not quantify Na_V_1.2 protein levels in the mice that underwent behavioral testing, precluding subject-level protein-phenotype correlations. Targeted Na_V_1.2 quantification in behavior-tested cohorts will be needed to assess causality. As with any mouse model, behavioral assays provide construct—not symptom—validity, and results should not be over-interpreted given the heterogeneity and multi-faceted nature of ASD and ID symptomology in humans. Finally, we behaviorally tested mice between 16-30 weeks of age, which is roughly equivalent to mid-adulthood in humans. Since *SCN2A* is important for early neuronal maturation^4^, future work should include observations earlier in development.

## Data Availability

The data that support the findings of this study are available on request from the corresponding authors.

## Acknowledgments

We thank Dr. Kevin Bender for providing training in whole-cell patch-clamp electrophysiology in acute mouse brain slices and Dr. Stephanie Gantz for hosting the experiments and providing access to—and assistance with—all slice preparation and recordings in her laboratory. We would like to thank the Williams and Ahern labs for critical feedback on this manuscript. We would also like to thank Katelin Scott and Julia Mullane for assistance with behavioral data scoring. We acknowledge the personnel and instrumentation in the Neural Circuits and Behavior Core in the Iowa Neuroscience Institute, supported in part by the Roy J. Carver Charitable Trust, UI Carver College of Medicine, and the Hawk-IDDRC (NIH P50HD103556).

