## Supplementary Information and Figures for "Position-dependent effects of *SCN2A* premature stop codons on neuronal excitability and behavior"

**Supplementary Material**

**Transgenic mouse generation**

Male mice older than 8 weeks were bred with 3 to 5-week-old super-ovulated females to produce zygotes for pronuclear injection. Female ICR (Envigo; Hsc:ICR(CD-1)) mice were used as recipients for embryo transfer.

For the *Scn2a^Y84X/+^* mice, chemically modified CRISPR-Cas12a crRNAs were purchased from IDT (Alt-R™ A.s. Cas12a crRNA). crRNA sequences were TTATTGATATAGTAGGGGTCCAG and TGAAAAGAGGCACCCTCTTCACCATCTTAAAGAAATCGAGGAAATAAAAGCAACTGGTCTAAAAACTCACCTTCTTATTGATGTATTATGGATCCAGGTCCTCCAGGGGCTCTGACACCATCTCTGGAGGAATGTCTCCATAAATAAAAGGAAGGGATTTTCCTGC. The crRNAs were suspended in T10E0.1 buffer at 1 μg/μl, heated at 98^o^C for 2 min, and allowed to cool slowly to 20^o^C in a thermal cycler. The annealed crRNAs were aliquoted into single-use tubes and stored at -80^o^C. Cas12a nuclease was also purchased from IDT (Alt-R™ A.s. Cas12a (Cpf1) V3; Catalog #1081068). Cas12a crRNA ribonucleoprotein complexes were made by combining Cas12a protein and crRNA in T10E0.1 (final concentrations: 1190 ng/μl (~7.3 μM) Cas12a protein and 760 ng/μl (~22.4 μM) crRNA). The Cas12a protein and annealed RNAs were incubated at 37^o^C for 10 min. The RNP complexes were combined with single-stranded oligonucleotide repair template and incubated an additional 5 min at 37^o^C. The final concentrations in the injection mix were 100 ng/μl (~0.6 μM) Cas12a protein, 64 ng/μl (~1.9 μM) crRNA, and 500 ng/μl oligonucleotide repair template. Pronuclear-stage embryos were collected using methods described elsewhere^1^. Embryos were collected in KSOM media (Millipore; MR101D) and washed 3 times to remove cumulus cells. Cas12a RNPs and single-stranded repair templates were injected^2^ into the pronuclei of the collected zygotes. Cas12a RNPs and oligonucleotide repair template were electroporated into zygotes briefly treated (10-sec) acid Tyrode’s solution (Sigma; T1788). The embryos were then washed 5 times with Opti-MEM, and 25-30 were aligned between the electrodes of a slide glass plate electrode with a 1.0 mm gap and platinum leads. The acid-treated embryos were overlaid with the electroporation mix and electroporated for 11 pulses at 30V (1-msec on:100-msec off). Electroporation was performed using an ECM830 square wave electroporator (BTX). Zygotes from both were incubated in KSOM with amino acids at 37^o^C under 5% CO_2_ until all zygotes were ready for injection. Fifteen to 25 embryos were immediately implanted into the oviducts of pseudo-pregnant ICR females.

For the *Scn2a^R1627X/+^* mice, chemically modified CRISPR-Cas9 crRNAs and CRISPR-Cas9 tracrRNA were purchased from IDT (Alt-R® CRISPR-Cas9 crRNA; Alt-R® CRISPR-Cas9 tracrRNA (Cat# 1072532)). crRNA sequences were Scn2a_R1627stop_5PB ATCTTACAAACTGCACATAT and Scn2a_R1627stop_3PD TTTTCAGGGTCACAATCGGG. The crRNAs and tracrRNA were suspended in T10E0.1 buffer and combined to 1 μg/μl (~29.5 μM) final concentration in a 1:2 (μg:μg) ratio. The RNAs were heated at 98^o^C for 2 min and allowed to cool slowly to 20^o^C in a thermal cycler. The annealed cr:tracrRNAs were aliquoted to single-use tubes and stored at -80^o^C. Cas9 nuclease was also purchased from IDT (Alt-R® S.p. HiFi Cas9 Nuclease). Cr:tracr:Cas9 ribonucleoprotein complexes were made by combining Cas9 protein and cr:tracrRNA in T10E0.1 (final concentrations: 300 ng/μl (~1.9 μM) Cas9 protein and 200 ng/μl (~5.9 μM) cr:tracrRNA). The Cas9 protein and annealed RNAs were incubated at 37^o^C for 10 min. The RNP complexes were combined with single-stranded repair template and incubated an additional 5 min at 37^o^C. The final concentrations in the injection mix were 60 ng/μl (~0.4 μM) Cas9 protein, 10 ng/μl (~0.3 μM) each cr:tracrRNA, and 20 ng/μl single-stranded repair template. Pronuclear-stage embryos were collected using methods described in^1^. Embryos were collected in KSOM media (Millipore; MR101D) and washed 3 times to remove cumulous cells. Cas9 RNPs and single-stranded repair template were injected^2^ into the pronuclei of the collected zygotes and incubated in KSOM with amino acids at 37^o^C under 5% CO_2_ until all zygotes were injected. Fifteen to 25 embryos were immediately implanted into the oviducts of pseudo-pregnant ICR females.

Both mouse lines were backcrossed onto the C57BL/6J background for at least four generations before incorporation into the study. To generate experimental animals, *Scn2a^Y84X/+^* or *Scn2a^R1627X/+^* mice were bred to WT mice to produce litters with combinations of *Scn2a^Y84X/+^* and WT offspring (or *Scn2a^R1627X/+^* and WT offspring, respectively). The lines were maintained separately, were not interbred, and transgenic mice were cohoused with their own WT littermates. Pups were weaned 21–24 days after birth.

**Genotyping**

Mice were genotyped at weaning via ear punch, and genotypes were confirmed via tail snips post-mortem. Genomic DNA (gDNA) was extracted with a tissue DNA lysate (NaOH and EDTA in ddH_2_O) and neutralization buffer (Tris-HCl in ddH_2_O). Polymerase chain reaction (PCR) was then performed on gDNA along with the following primers: Y84X Scn2a_Tyr84TAA:F:5′GAACAACAGGCTCAACCAAATG3′ and Scn2a_Tyr84TAA:R:5′GGGAATCCCTTGCTGCTATT3′; or R1627X LG366_R1627:F:5’CCCACCCTCTGATGGTCTTA3’ and LG236_Scn2a_R1627:R:5’ACCACAACCAGGAAGGATATG3’), GoTaq(R) G2 Green Master Mix (Catalog #M7823, Promega, Madison, WI), and ddH_2_O. PCR conditions for the Y84X line were 95^o^C for 2 min, then 35 cycles of 95^o^C for 25 sec, 62^o^C for 25 sec, and 72^o^C for 1 min 15 sec, and a final inactivation step of 72^o^C for 5 min. For the R1627X line, PCR conditions were 95^o^C for 2 min, then 37 cycles of 98^o^C for 20 sec, 61^o^C for 30 sec, and 72^o^C for 2 min, and a final inactivation step of 72^o^C for 7 min. The PCR product was then digested with a 1:1 ratio of BamHI-HF and rCutSmart buffer for the Y84X line (Catalog #R3136S, New England Biolabs, Ipswich, MA), or a 1:1 ratio of DdeI and rCutSmart buffer for the R1627X line (Catalog #R0175S, New England Biolabs) at 37^o^C for 45 min. Following enzyme restriction, the samples were separated by electrophoresis on a 2% agarose gel.

**Sanger sequencing** **Primers**

*Scn2a^Y84X/+^* mouse samples: AAATGGCCCAAAGCCAAATAG

*Scn2a^R1627X/+^* mouse samples: CATCTCAAAGTCGTCCTCACTC

**PCR amplification Primers**

*Scn2a^Y84X/+^* mouse samples:

Forward Primer: CAAGGACGAAGACGACGAAA

Reverse Primer: GGCAGTACCATTCCAATCCA

*Scn2a^R1627X/+^* mouse samples:

Forward Primer: GCGGAGCTGATAGAGAAGTATT

Reverse Primer 1: CGCTTCAGAGTGGTGGTAAT

Reverse Primer 2: CATAAGAGACCTTGGAGGGATTG

**General behavioral procedures**

All mice were adults (16-30 weeks) at the time of behavioral testing. Sample sizes are indicated in each figure. Behavioral experiments were run in two independent cohorts of mice in the following order: Cohort 1 underwent elevated zero maze, open field test, rotarod, and electroencephalogram (EEG) and electromyography (EMG) monitoring; Cohort 2 underwent three-chamber social task, prepulse inhibition, and maximal electroshock (MES) seizure induction. A minimum of 2 days was taken between every behavioral assay to reduce the impact of stress from previous tests on subsequent tests. Mice were randomized and color-coded with Sharpies on their tails at the beginning of behavioral testing to maintain blinding during testing.

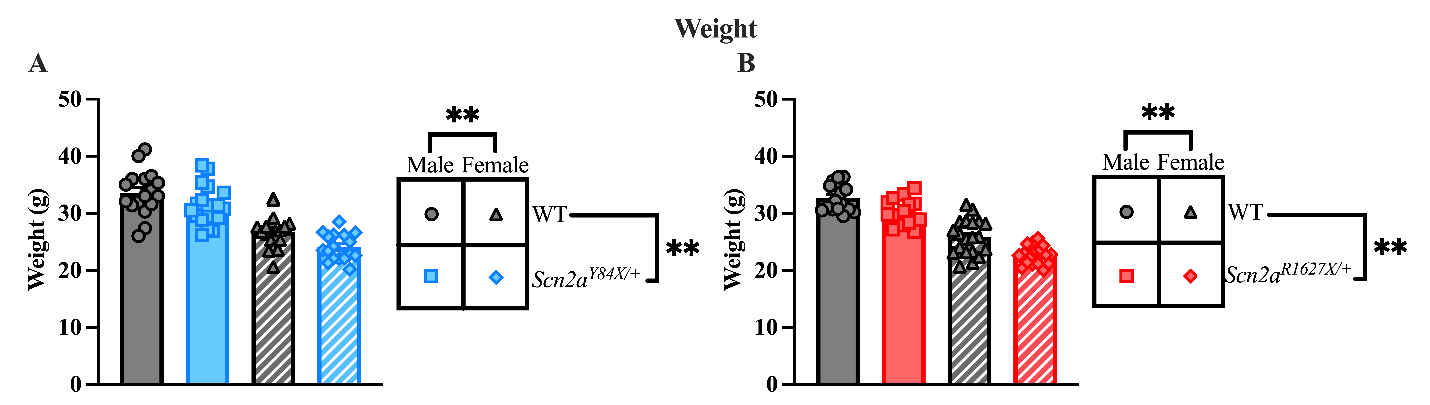

**Supplementary Figure 1. *Scn2a^Y84X/+^* and *Scn2a^R1627X/+^* mice are on average smaller than their WT littermates.** (**A)** In the Y84X line, *Scn2a^Y84X/+^* mice are on average 2.03 grams smaller than WT mice. Females were on average 6.81 grams smaller than males. **(B)** In the R1627X line, *Scn2a^R1627X/+^* mice are on average 2.31 grams smaller than WT mice. Females were on average 7.06 grams smaller than males. **(A-B)** Data are expressed as mean ± SEM. **(A)** *n* = 73 (WTs: *n* = 17 males, *n* = 19 females; *Scn2a^Y84X/+^*: *n* = 20 males, *n* = 17 females). **(B)** *n* = 69 (WTs: *n* = 16 males, *n* = 20 females; *Scn2a^R1627X/+^*: *n* = 16 males, *n* = 17 females).

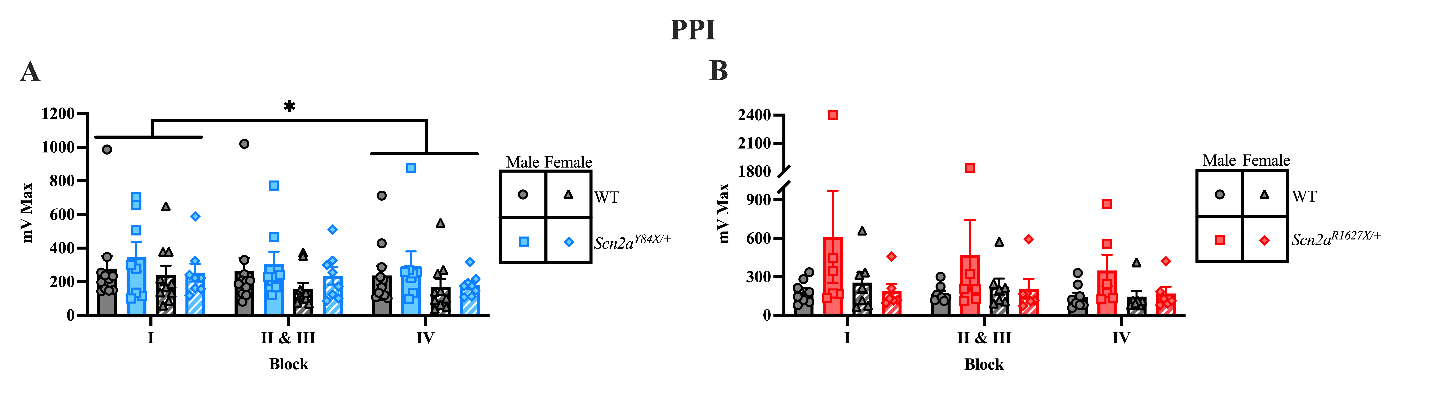

**Supplementary Figure 2. There is a mild habituation to the auditory startle. (A)** All mice in the Y84X line had a decrease in startle response over the course of the experiment. **(B)** There were no differences in the auditory startle response in the R1627X line. **(A-B)** Data are expressed as mean ± SEM. **(A)** *n* = 27 (WTs: *n* = 8 males, *n* = 7 females; *Scn2a^R1627X/+^*: *n* = 6 males, *n* = 6 females). **(B)** *n* = 37 (WTs: *n* = 11 males, *n* = 10 females; *Scn2a^Y84X/+^*: *n* = 8 males, *n* = 8 females).

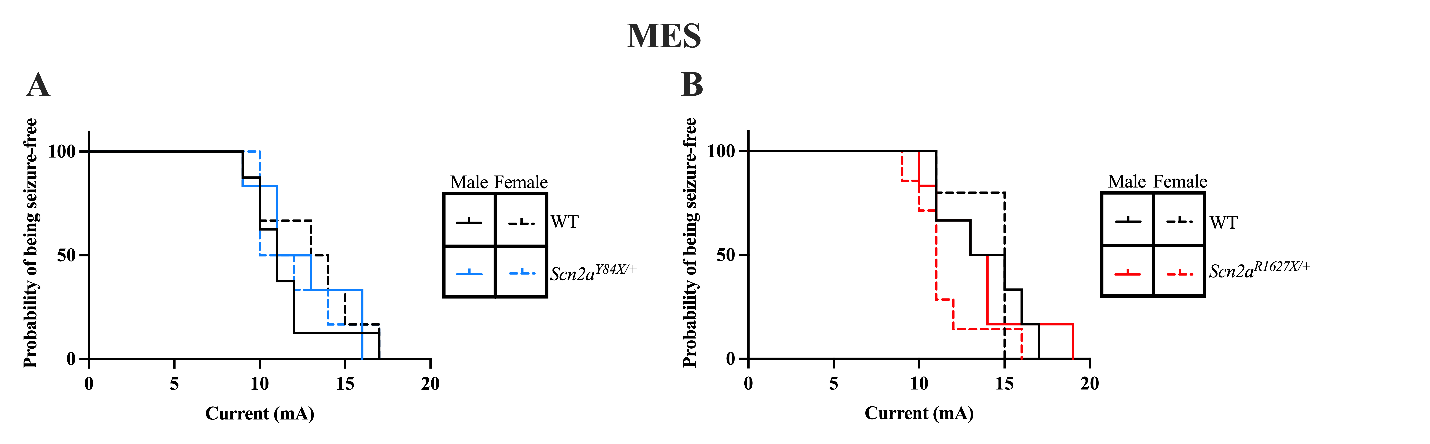

**Supplementary Figure 3. No differences in the probability of being seizure-free.** **(A)** There were no differences observed in probability of being seizure-free in the Y84X line or **(B)** the R1627X line. **(A-B)** Data are expressed as mean ± SEM. **(A)** *n* = 26 (WTs: *n* = 8 males, *n* = 6 females; *Scn2a^Y84X/+^*: *n* = 6 males, *n* = 6 females). **(B)** *n* = 24 (WTs: *n* = 6 males, *n* = 5 females; *Scn2a^R1627X/+^*: *n* = 6 males, *n* = 7 females).

**Supplementary Table 1. Effects of genotype and sex on weight in Y84X and R1627X mice**

| **Mouse line** | **Effect** | **NumDF, DenDF** | **F-value** | ***P*-value** |
| --- | --- | --- | --- | --- |
| **Y84X** | Genotype | 1, 69 | 11.39 | **< 0.01** |
|  | Sex | 1, 69 | 87.87 | **< 0.01** |
|  | Genotype × Sex | 1, 69 | 0.067 | 0.796 |
| **R1627X** | Genotype | 1, 65 | 23.83 | **< 0.01** |
|  | Sex | 1, 65 | 150.0 | **< 0.01** |
|  | Genotype × Sex | 1, 65 | 0.289 | 0.593 |

Two-way ANOVA results for effects of genotype, sex, and genotype × sex interaction on body weight in Y84X and R1627X lines. Bold values indicate statistically significant effects (*p* < 0.05).

**Supplementary Table 2. Effects of genotype and sex on distance traveled in the elevated zero maze (EZM) and open field test (OFT)**

| **Task** | **Mouse line** | **Measure** | **Effect** | **NumDF, DenDF** | **F-value** | ***P*-value** |
| --- | --- | --- | --- | --- | --- | --- |
| **EZM** | Y84X | Distance traveled | Genotype | 1, 29 | 1.686 | 0.204 |
|  |  |  | Sex | 1, 29 | 0.091 | 0.765 |
|  |  |  | Genotype × Sex | 1, 29 | 2.252 | 0.144 |
|  | R1627X |  | Genotype | 1, 37 | 0.638 | 0.429 |
|  |  |  | Sex | 1, 37 | 7.007 | **0.012** |
|  |  |  | Genotype × Sex | 1, 37 | 0.239 | 0.628 |
| **OFT** | Y84X | Distance traveled | Genotype | 1, 32 | 0.424 | 0.519 |
|  |  |  | Sex | 1, 32 | 0.003 | 0.958 |
|  |  |  | Genotype × Sex | 1, 32 | 0.002 | 0.961 |
|  | R1627X |  | Genotype | 1, 38 | 3.119 | 0.085 |
|  |  |  | Sex | 1, 38 | 0.559 | 0.459 |
|  |  |  | Genotype × Sex | 1, 38 | 0.038 | 0.846 |

Two-way ANOVA results for effects of genotype, sex, and genotype × sex interaction on distance traveled in the EZM and OFT. Bold values indicate statistically significant effects (*p* < 0.05).

**Supplementary Table 3. Effects of genotype and sex in learning the accelerated rotarod task**

| **Mouse line** | **Effect** | **NumDF, DenDF** | **F-value** | ***P*-value** |
| --- | --- | --- | --- | --- |
| **Y84X** | Genotype | 1, 32 | 1.47 | 0.234 |
|  | Sex | 1, 32 | 3.609 | 0.067 |
|  | Session | 4, 128 | 54.55 | **< 0.01** |
|  | Genotype × Sex | 1, 32 | 1.495 | 0.230 |
|  | Genotype × Session | 4, 128 | 0.636 | 0.638 |
|  | Sex × Session^a^ | 4, 128 | 4.189 | **< 0.01** |
|  | Genotype × Sex × Session^b^ | 4, 128 | 4.639 | **< 0.01** |
| **R1627X** | Genotype | 1, 38 | 2.289 | 0.139 |
|  | Sex | 1, 38 | 0.222 | 0.640 |
|  | Session | 4, 152 | 62.63 | **< 0.01** |
|  | Genotype × Sex | 1, 38 | 0.118 | 0.733 |
|  | Genotype × Session | 4, 152 | 1.125 | 0.347 |
|  | Sex × Session | 4, 152 | 0.868 | 0.485 |
|  | Genotype × Sex × Session | 4, 152 | 0.426 | 0.789 |

Linear-mixed effects model results for effects of genotype, sex, and interactions in latency to fall on the accelerated rotarod task. Bold values indicate statistically significant effects (*p* < 0.05). Post hoc tests were conducted following significant interactions using estimated marginal means.

^a^Post hoc comparisons were conducted to follow-up the significant sex × session interaction in the Y84X line. We observed a significant difference between males and females on session 5 (session 1, t_69.8_ = 0.165, *p* = 0.869, session 2, t_69.8_ = 1.082, *p* = 0.283, session 3, t_69.8_ = 0.830, *p* = 0.409, session 4, t_69.8_ = 1.960, *p* = 0.054, session 5, t_69.8_ = 3.649, *p* < 0.01).

^b^Post hoc comparisons were conducted to follow-up the significant genotype × sex × session interaction in the Y84X line. Among females, no significant differences were observed between *Scn2a^Y84X/+^* and WT mice (session 1, t_69.8_ = 0.561, *p* = 0.577, session 2, t_69.8_ = 1.848, *p* = 0.069, session 3, t_69.8_ = -1.068, *p* = 0.289, session 4, t_69.8_ = -1.158, *p* = 0.251, session 5, t_69.8_ = -0.214, *p* = 0.832). In males, there was a significant difference between *Scn2a^Y84X/+^* and WT mice on sessions 3 and 4, which resolved on session 5 (session 1, t_69.8_ = 1.008, *p* = 0.317, session 2, t_69.8_ = 0.624, *p* = 0.535, session 3, t_69.8_ = 2.179, *p* = 0.033, session 4, t_69.8_ = 2.394, *p* = 0.019, session 5, t_69.8_ = 0.567, *p* = 0.573).

**Supplementary Table 4. Effects of genotype and sex on anxiety-like behavior in the elevated zero maze (EZM) and open field test (OFT)**

| **Task** | **Mouse line** | **Measure** | **Effect** | **NumDF, DenDF** | **F-value** | ***P*-value** |
| --- | --- | --- | --- | --- | --- | --- |
| **EZM** | Y84X | Time in open area | Genotype | 1, 29 | 9.541 | **< 0.01** |
|  |  |  | Sex | 1, 29 | 1.916 | 0.177 |
|  |  |  | Genotype × Sex | 1, 29 | 1.132 | 0.296 |
|  | R1627X |  | Genotype | 1, 37 | 0.175 | 0.678 |
|  |  |  | Sex | 1, 37 | 0.563 | 0.458 |
|  |  |  | Genotype × Sex | 1, 37 | 0.187 | 0.668 |
| **OFT** | Y84X | Time in inner area | Genotype | 1, 32 | 0.000 | 0.987 |
|  |  |  | Sex | 1, 32 | 1.447 | 0.238 |
|  |  |  | Genotype × Sex | 1, 32 | 0.748 | 0.394 |
|  | R1627X |  | Genotype | 1, 38 | 3.010 | 0.091 |
|  |  |  | Sex | 1, 38 | 0.062 | 0.805 |
|  |  |  | Genotype × Sex | 1, 38 | 1.545 | 0.221 |

Two-way ANOVA and ranked two-way ANOVA results for effects of genotype, sex, and genotype × sex interaction on anxiety-like behavior in the EZM and OFT. Bold values indicate statistically significant effects (*p* < 0.05).

**Supplementary Table 5. Effects of genotype and sex on grooming behavior in the open field test (OFT)**

| **Mouse line** | **Measure** | **Effect** | **NumDF, DenDF** | **F-value** | ***P*-value** |
| --- | --- | --- | --- | --- | --- |
| Y84X | Grooming frequency | Genotype | 1, 32 | 4.018 | 0.054 |
|  |  | Sex | 1, 32 | 2.081 | 0.159 |
|  |  | Genotype × Sex | 1, 32 | 0.009 | 0.926 |
| R1627X |  | Genotype | 1, 38 | 11.93 | **< 0.01** |
|  |  | Sex | 1, 38 | 0.030 | 0.864 |
|  |  | Genotype × Sex^a^ | 1, 38 | 4.216 | **0.047** |
| Y84X | Time spent grooming | Genotype | 1, 32 | 5.071 | **0.031** |
|  |  | Sex | 1, 32 | 0.702 | 0.408 |
|  |  | Genotype × Sex | 1, 32 | 1.243 | 0.273 |
| R1627X |  | Genotype | 1, 38 | 11.39 | **< 0.01** |
|  |  | Sex | 1, 38 | 0.029 | 0.865 |
|  |  | Genotype × Sex | 1, 38 | 0.479 | 0.493 |

Two-way ANOVA and ranked two-way ANOVA results for effects of genotype, sex, and genotype × sex interaction on grooming behaviors in the OFT. Bold values indicate statistically significant effects (*p* < 0.05). Post hoc tests were conducted following significant interactions using uncorrected Fisher’s LSD.

^a^Post hoc comparisons were conducted to follow-up the significant genotype × sex interaction in the R1627X line. Among females, no significant differences were observed between *Scn2a^R1627X/+^* and WT mice (t_38_ = 0.998, *p* = 0.325). In males, there was a significant difference between *Scn2a^R1627X/+^* and WT mice (t_38_ = 3.475, *p* < 0.01).

**Supplementary Table 6. Effects of genotype and sex on distance traveled and sociability in the three-chamber social task (3-CST)**

**(A) Group effects: linear-mixed effects model on distance traveled and sociability**

| **Mouse line** | **Measure** | **Effect** | **NumDF, DenDF** | **F-value** | ***P*-value** |
| --- | --- | --- | --- | --- | --- |
| Y84X | Distance traveled in habituation and sociability phases | Genotype | 1, 32.44 | 0.132 | 0.718 |
|  |  | Sex | 1, 32.44 | 0.313 | 0.579 |
|  |  | Phase | 1, 30.88 | 174.278 | **< 0.01** |
|  |  | Genotype × Sex | 1, 32.44 | 1.724 | 0.198 |
|  |  | Genotype × Phase | 1, 30.88 | 2.009 | 0.167 |
|  |  | Sex × Phase | 1, 30.88 | 2.682 | 0.112 |
|  |  | Genotype × Sex × Phase | 1, 30.88 | 0.012 | 0.915 |
| R1627X |  | Genotype | 1, 23 | 0.974 | 0.334 |
|  |  | Sex | 1, 23 | 3.156 | 0.088 |
|  |  | Phase | 1, 23 | 73.67 | **< 0.01** |
|  |  | Genotype × Sex | 1, 23 | 2.489 | 0.128 |
|  |  | Genotype × Phase | 1, 23 | 0.095 | 0.760 |
|  |  | Sex × Phase | 1, 23 | 0.395 | 0.536 |
|  |  | Genotype × Sex × Phase | 1, 23 | 3.327 | 0.081 |
| Y84X | Social preference in sociability phase | Genotype | 1, 31 | 0.000 | 0.999 |
|  |  | Sex | 1, 31 | 0.151 | 0.701 |
|  |  | Cylinder | 1, 31 | 42.21 | **< 0.01** |
|  |  | Genotype × Sex | 1, 31 | 0.044 | 0.835 |
|  |  | Genotype × Cylinder | 1, 31 | 0.193 | 0.664 |
|  |  | Sex × Cylinder | 1, 31 | 0.212 | 0.649 |
|  |  | Genotype × Sex × Cylinder | 1, 31 | 1.471 | 0.234 |
| R1627X |  | Genotype | 1, 46 | 0.761 | 0.388 |
|  |  | Sex | 1, 46 | 0.438 | 0.511 |
|  |  | Cylinder | 1, 46 | 65.32 | **< 0.01** |
|  |  | Genotype × Sex | 1, 46 | 0.793 | 0.378 |
|  |  | Genotype × Cylinder | 1, 46 | 0.030 | 0.863 |
|  |  | Sex × Cylinder | 1, 46 | 0.016 | 0.901 |
|  |  | Genotype × Sex × Cylinder | 1, 46 | 0.734 | 0.396 |

**(B) Social preference: paired t-test within groups**

| **Mouse line** | **Measure** | **Sex and genotype** | **t(DF)** | **P-value** |
| --- | --- | --- | --- | --- |
| Y84X | Social vs object cylinder | Male WT | t(10) = 5.860 | **< 0.01** |
|  |  | Male *Scn2a^Y84X/+^* | t(5) = 3.710 | **0.014** |
|  |  | Female WT | t(9) = 1.865 | 0.095 |
|  |  | Female *Scn2a^Y84X/+^* | t(8) = 3.60 | **< 0.01** |
| R1627X |  | Male WT | t(7) = 3.657 | **< 0.01** |
|  |  | Male *Scn2a^R1627X/+^* | t(5) = 5.389 | **< 0.01** |
|  |  | Female WT | t(6) = 5.622 | **< 0.01** |
|  |  | Female *Scn2a^R1627X/+^* | t(5) = 3.120 | **0.026** |

Linear mixed effects model and linear model used to assess the effects of genotype and sex on distance traveled and time spent with each cylinder in the sociability phase of the 3-CST. Paired t-test results to compare time within-group preference for the social versus object stimuli. Bold values indicate statistically significant effects (*p* < 0.05).

**Supplementary Table 7. Effects of genotype and sex on sensorimotor gating and startle response during the prepulse inhibition task (PPI)**

| **Mouse line** | **Measure** | **Effect** | **NumDF, DenDF** | **F-value** | ***P*-value** |
| --- | --- | --- | --- | --- | --- |
| Y84X | Percent of prepulse inhibition | Genotype | 1, 33 | 1.687 | 0.203 |
|  |  | Sex | 1, 33 | 2.843 | 0.101 |
|  |  | Decibel | 2, 66 | 73.88 | **< 0.01** |
|  |  | Genotype × Sex | 1, 33 | 1.00 | 0.325 |
|  |  | Genotype × Decibel | 2, 66 | 0.296 | 0.745 |
|  |  | Sex × Decibel | 2, 66 | 0.189 | 0.828 |
|  |  | Genotype × Sex × Decibel | 2, 66 | 1.421 | 0.249 |
| R1627X |  | Genotype | 1, 23 | 0.421 | 0.523 |
|  |  | Sex | 1, 23 | 0.559 | 0.461 |
|  |  | Decibel | 2, 46 | 66.49 | **< 0.01** |
|  |  | Genotype × Sex | 1, 23 | 0.835 | 0.370 |
|  |  | Genotype × Decibel | 2, 46 | 0.029 | 0.971 |
|  |  | Sex × Decibel | 2, 46 | 0.188 | 0.829 |
|  |  | Genotype × Sex × Decibel | 2, 46 | 0.416 | 0.662 |
| Y84X | Startle response | Genotype | 1, 33 | 0.570 | 0.456 |
|  |  | Sex | 1, 33 | 1.888 | 0.179 |
|  |  | Block | 2, 66 | 4.453 | **0.015** |
|  |  | Genotype × Sex | 1, 33 | 0.033 | 0.857 |
|  |  | Genotype × Block | 2, 66 | 0.170 | 0.844 |
|  |  | Sex × Block | 2, 66 | 0.191 | 0.827 |
|  |  | Genotype × Sex × Block | 2, 66 | 0.749 | 0.477 |
| R1627X |  | Genotype | 1, 23 | 1.515 | 0.231 |
|  |  | Sex | 1, 23 | 1.069 | 0.312 |
|  |  | Block | 2, 46 | 3.028 | 0.058 |
|  |  | Genotype × Sex | 1, 23 | 1.938 | 0.177 |
|  |  | Genotype × Block | 2, 46 | 0.283 | 0.755 |
|  |  | Sex × Block | 2, 46 | 0.547 | 0.583 |
|  |  | Genotype × Sex × Block | 2, 46 | 1.623 | 0.208 |

Linear-mixed effects model results for effects of genotype, sex, and interactions in percent of prepulse inhibition and startle response in PPI task. Bold values indicate statistically significant effects (*p* < 0.05).

**Supplementary Table 8. Effects of genotype and sex on mortality, current at which seizures were induced, and extension-to-flexion (E/F) ratio in the maximal electroshock (MES) assay**

| **Mouse line** | **Measure** | **Effect** | **Statistic** | **DF** | **Effect size/CI** | ***P*-value** |
| --- | --- | --- | --- | --- | --- | --- |
| Y84X | Mortality | Genotype | OR = 13.14 | – | CI [1.268, 704.98] | **0.014** |
|  |  | Sex | OR = 3.476 | – | CI [0.513, 29.87] | 0.217 |
| R1627X |  | Genotype | 𝜒^2^ = 4.086 | 1 | Cramer’s V = 0.497 | **0.043** |
|  |  | Sex | 𝜒^2^ = 0.671 | 1 | Cramer’s V = 0.251 | 0.413 |
| Y84X | Current at which seizures were induced | Genotype | F = 0.050 | 1, 22 | – | 0.825 |
|  |  | Sex | F = 0.166 | 1, 22 | – | 0.688 |
|  |  | Genotype × Sex | F = 0.649 | 1, 22 | – | 0.429 |
| R1627X |  | Genotype | F = 3.508 | 1, 20 | – | 0.076 |
|  |  | Sex | F = 0.494 | 1, 20 | – | 0.490 |
|  |  | Genotype × Sex | F = 0.942 | 1, 20 | – | 0.343 |
| Y84X | E/F ratio | Genotype | F = 0.324 | 1, 20 | – | 0.575 |
|  |  | Sex | F = 0.029 | 1, 20 | – | 0.866 |
|  |  | Genotype × Sex | F = 2.027 | 1, 20 | – | 0.170 |
| R1627X |  | Genotype | F = 2.059 | 1, 20 | – | 0.167 |
|  |  | Sex | F = 0.359 | 1, 20 | – | 0.555 |
|  |  | Genotype × Sex | F = 0.639 | 1, 20 | – | 0.434 |
| Y84X | Chance of being-seizure free | Genotype | 𝜒^2^ = 0.000 | 1 | – | 0.900 |
|  |  | Sex | 𝜒^2^ = 0.300 | 1 | – | 0.600 |
| R1627X |  | Genotype | 𝜒^2^ = 1.600 | 1 | – | 0.200 |
|  |  | Sex | 𝜒^2^ = 0.800 | 1 | – | 0.400 |

Fisher’s exact test, Chi-square test of independence, ranked two-way ANOVA, two-way ANOVA, and log-rank test results for effects of genotype, sex, and interactions in mortality, current at which seizures were induced, E/F ratio, and chance of being seizure-free in the MES threshold assay. Bold values indicate statistically significant effects (*p* < 0.05).
